# An unbiased screening approach for identifying proteins with a mechanosensitive nuclear localization

**DOI:** 10.1101/2025.09.17.676718

**Authors:** Pei-Li Tseng, Weiwei Sun, Jiawei Li, Mark O. Collins, Kai S. Erdmann

**Affiliations:** University of Sheffield

## Abstract

Mechanical forces profoundly influence a wide range of cellular behaviors. However, the mechanisms by which these forces are sensed and transduced into specific biological signals remain incompletely understood. One established principle of mechanotransduction is the mechanical regulation of subcellular protein localization, which enables control over the activity of key regulators such as transcription and splicing factors. In this protocol, we present a screening approach to identify proteins that exhibit mechanosensitive nuclear localization. We describe the engineering of cell lines with tuneable actomyosin contractility and the application of a spatially restricted in situ biotinylation strategy. This method, combined with liquid chromatography-tandem mass spectrometry (LC-MS/MS), enables the identification of proteins whose nuclear localization changes in response to mechanical cues. For a detailed application of this protocol, please refer to Tseng et al. (2025) [1]

## BEFORE YOU BEGIN

Cells sense and respond to mechanical stimuli from their extracellular microenvironment through mechanotransduction, a process that converts physical forces into biochemical signals to regulate cell behaviour, tissue homeostasis and disease progression [2-4]. A central player of this process is the small GTPase RhoA, which is activated by external mechanical cues such as extracellular matrix stiffening and tension at adherens junctions [5]. Upon activation, RhoA initiates a signaling cascade that drives actomyosin contractility and actin polymerization, dynamically reshaping the cytoskeleton to mediate cellular responses to force [6, 7].

Mechanical forces are transmitted to the nucleus via the Linker of Nucleoskeleton and Cytoskeleton (LINC) complex [8]. This transmission leads to nuclear membrane deformation and modulates the architecture and function of nuclear pore complex [9], ultimately affecting nuclear transport, chromatin organization, and transcriptional regulation [10-12]. These observations support the nucleus’s dual role as both a mechanical sensor and regulatory hub, highlighting the importance of identifying proteins with a mechanosensitive nuclear localisation and to better understand their roles in mechanotransduction.

Physical methods to isolate intact nuclei for biochemical analysis have the potential to interfere with protein changes relevant to mechanotransduction. To circumvent this problem, we implemented nuclear-restricted biotinylation of cells in situ, allowing for the purification of nuclear proteins from cell lysates. Here we present a protocol to systematically identify proteins exhibiting mechanosensitive nuclear localization using a TurboID-based proximity labeling approach combined with label-free mass spectrometry. We utilize a HEK293-tet-RhoA-TurboID cell model, in which constitutively active RhoA is expressed under the control of a tetracycline-inducible promoter. Activation of RhoA mimics mechanical stimulation and triggers nuclear responses. The TurboID enzyme, fused to a nuclear localization signal, enables selective biotinylation of nuclear proteins in living cells, which can then be isolated and analysed by mass spectrometry-based proteomics.

The protocol consists of three main parts: (1) generation of a cell model with tuneable actomyosin contractility for screening proteins with a mechanosensitive nuclear localization, (2) quantitative proteomic analysis of biotinylated proteins by LC-MS/MS, and (3) validation of mechanosensitive subcellular localisation of candidate proteins. We use YAP, a well-characterized mechanosensitive transcriptional regulator whose nuclear localization is modulated by mechanical stimuli [13, 14], to validate our screen approach.

### Construction of pcDNA5/FRT/TO-RhoA-Q63L

Timing: 1 week

To identify proteins that change nuclear abundance in response to mechanical stimuli, we employ an approach that mimics mechanosensitive signalling initiated by RhoA. In this protocol, we engineer a plasmid encoding an EGFP-tagged, constitutively active RhoA variant (RhoA-Q63L) under a tetracycline inducible promoter. Using standard cloning techniques, we isolate EGFP-RhoA-Q63L from donor plasmid pcDNA3-EGFP-RhoA-Q63L and subclone it into the recipient plasmid pcDNA5/FRT/TO to generate pcDNA5/FRT/TO-RhoA-Q63L.

### Construction of pcDNA3.1-Puro-3xHA-TurboID-NLS

Timing: 1 week

TurboID is an engineered biotin ligase variant with enhanced biotinylation kinetics, capable of rapid labelling of proximal proteins within a ∼10 nm radius [15, 16]. To biotinylate selective nuclear proteins, we utilise a plasmid coding for an HA-tagged TurboID including three nuclear localization signals (NLS) for nuclear-restricted TurboID expression (Figure 1). Using standard cloning techniques, we isolate 3xHA-TurboID-NLS from the donor plasmid 3xHA-TurboID-NLS_pCDNA3 and insert into the recipient vector pCDNA3.1-PuroR to create recombinant plasmid pcDNA3.1-Puro-3xHA-TurboID-NLS. This construct enables nuclear specific protein labelling and puromycin selection of transfected cells.

**Figure 1.**
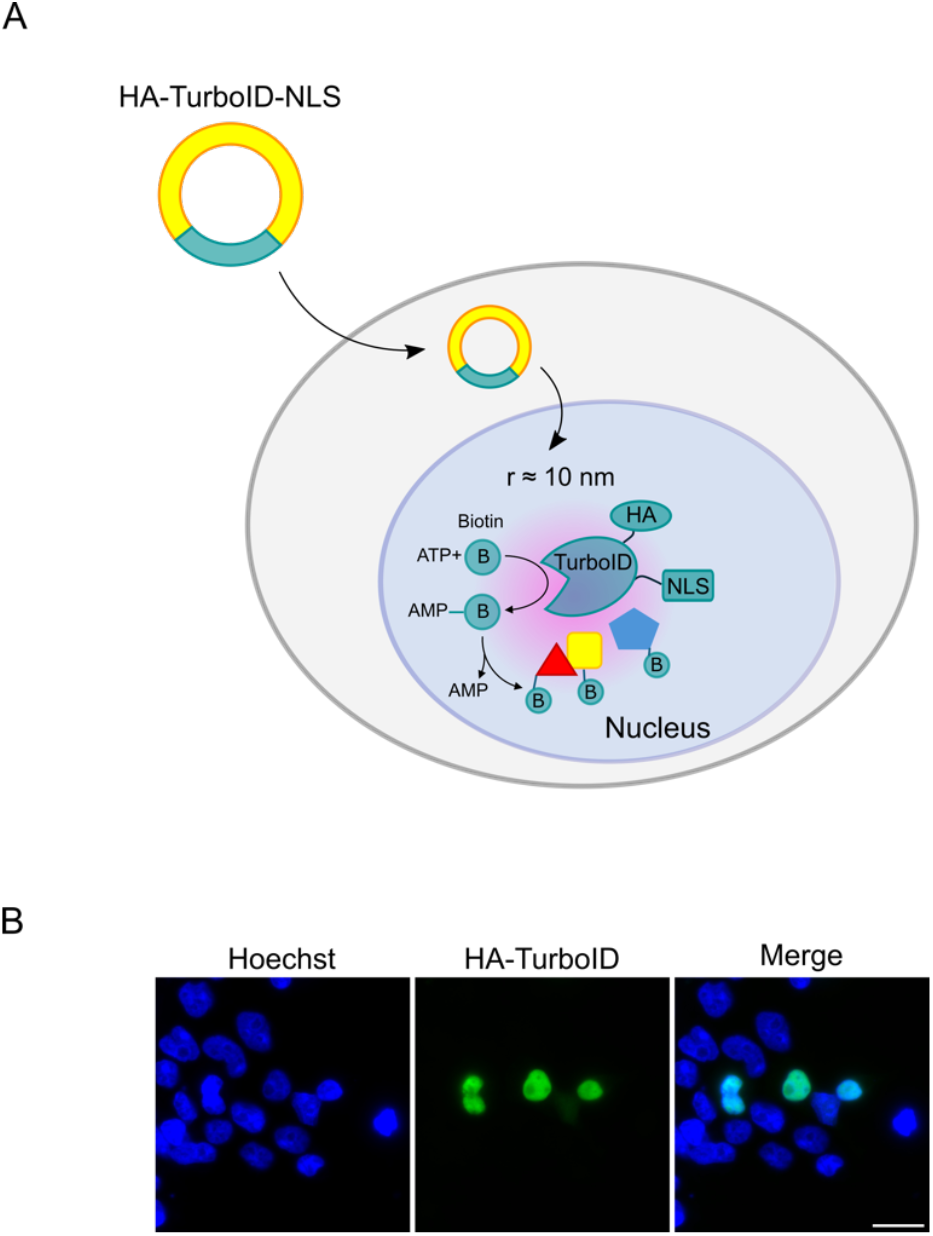
Nuclear localisation of TurboID-NLS. **A**: Schematic representation of 3xHA-TurboID-NLS mediated proximity biotinylation. **B**: Immunofluorescence staining confirms nuclear localisation of 3xHA-TurboID-NLS in transiently transfected HEK293 cells. Bar represents 50 μm.

***Note*:** The 3xHA-TurboID-NLS_pCDNA3 (Addgene_107171) contains a *Neo/KanR* for selection. However, our kill curve test shows that HEK293-tet-RhoA is weakly sensitive to geneticin, which hinders the selection of stable clones. Thus, we re-cloned 3xHA-TurboID-NLS into another recipient vector pCDNA3.1-PuroR to generate pcDNA3.1-Puro-3xHA-TurboID-NLS. If your cells are sensitive to geneticin, you can skip this step.

## KEY RESOURCES TABLE

| REAGENT or RESOURCE | SOURCE | IDENTIFIER |
| --- | --- | --- |
| Antibodies |  |  |
| Mouse monoclonal anti-YAP | Santa Cruz | #sc-101199; RRID: AB_1131430 |
| Rabbit polyclonal anti-GFP | Invitrogen | #A-11122; RRID: AB_221569 |
| Mouse monoclonal anti-nucleolin | Santa Cruz | #sc-8031; RRID: AB_670271 |
| Mouse monoclonal anti-Integrin β1 | Santa Cruz | #sc-374429; RRID: AB_11012020 |
| Mouse monoclonal anti-γ-adaptin | Lab of Professor Andrew Peden | N/A |
| Rabbit monoclonal anti-HA tag | Cell Signalling | #3724; RRID: AB_1549585 |
| Mouse monoclonal anti-PTBP1 | Thermo Fisher Scientific | #32-4800; RRID: AB_2533082 |
| Alexa Fluor™ 680 Streptavidin Conjugate | Thermo Fisher Scientific | #S32358 |
| Donkey anti-mouse IgG (H+L) Alexa Fluor™ 594 | Thermo Fisher Scientific | #A-11005; RRID: AB_2534073 |
| Goat anti-mouse IgG (H+L) Alexa Fluor™ 488 | Thermo Fisher Scientific | #A-11001; RRID: AB_2534069 |
| DyLight™800 4X PEG conjugate anti- mouse IgG (H+L) | Thermo Fisher Scientific | #SA5-35521; RRID: AB_2556774 |
| Experimental models: Cell lines |  |  |
| Flp-In™ T-REX™ HEK293 | Thermo Fisher Scientific | RRID: CVCL_U427 |
| MCF10A | ATCC | #CRL-10317; RRID: CVCL_0598 |
| NIH3T3 | ATCC | #CRL-1658; RRID: CVCL_0594 |
| HEK293-tet-RhoA-TurboID | This study | N/A |
| Chemicals, peptides, and recombinant proteins |  |  |
| DMEM-GlutaMax | Gibco | #10569010 |
| Blasticidin S HCl | Gibco | #R21001 |
| Zeocin™ Selection Reagent | Gibco | #R25001 |
| Hygromycin B | Gibco | #10687010 |
| Puromycin dihydrochloride | Santa Cruz | #sc-108071 |
| Tetracycline hydrochloride | Sigma-Aldrich | #T7660 |
| Biotin | Sigma-Aldrich | #B4501 |
| Collagen I, rat tail | Gibco | #A1048301 |
| Hoechst 33342, trihydrochloride | Thermo Fisher Scientific | #H3570 |
| ML-7, hydrochloride | Sigma-Aldrich | #475880 |
| ProLong™ Gold Antifade Mountant | Invitrogen | #P36930 |
| Dimethyl Sulfoxide Hybri-Max™ Sterile-Filtered BioReagent (DMSO) | Sigma-Aldrich | #D2650-100ML |
| Pierce™ Streptavidin Agarose | Thermo Fisher Scientific | #20349 |
| Lipofectamine™ 2000 | Thermo Fisher Scientific | #11668019 |
| cOmplete™, EDTA-free Protease Inhibitor Cocktail | Sigma-Aldrich | #11873580001 |
| Ammonium bicarbonate | Thermo Fisher Scientific | #393212500 |
| TCEP (Tris(2-carboxyethyl)phosphine) | Sigma-Aldrich | #C4706 |
| IAA (Iodoacetamide) | Sigma-Aldrich | #I1149 |
| TFA (Trifluoroacetic acid) | Sigma-Aldrich | #T6508 |
| ACN (Acetonitrile) | Sigma-Aldrich | #34851 |
| Water, for HPLC-MS, Fisher Chemical™ | Fisher Scientific | #10777404 |
| Deposited data |  |  |
| Proteomics Data |  | PXD046157 |
| Recombinant DNA |  |  |
| pcDNA5/FRT/TO | Thermo Fisher Scientific | #V652020 |
| pOG44 | Thermo Fisher Scientific | #V600520 |
| pcDNA3-EGFP-RhoA-Q63L | Addgene | RRID:<br>Addgene_12968 |
| 3xHA-TurboID-NLS_pCDNA3 | Addgene | RRID:<br>Addgene_107171 |
| pcDNA3.1 (+)-myc-PuroR | Lab of Dr Kai Erdmann | N/A |
| Software and algorithms |  |  |
| MaxQuant 1.6.2.6 | MaxQuant | RRID:SCR_014485 |
| Perseus 1.5.6.0 | MaxQuant | RRID:SCR_015753 |
| Fiji | (Schindelin et al. 2012) | RRID:SCR_002285 |
| ImageStudioLite | LI-COR Biosciences | N/A |
| GraphPad Prism 9.5.1 | Prism | RRID:SCR_002798 |
| Biorender | Biorender | RRID:SCR_018361 |
| Other |  |  |
| Pierce® Streptavidin Agarose Resins | Thermo Fisher Scientific | #20349 |
| Scienceware® cloning discs | Sigma-Aldrich | Discontinued |
| Pierce™ C18 stage tips | Thermo Fisher Scientific | #87782 |
| Wizard® Minicolumn | Promega | #A7211 |
| Orbitrap Elite Hybrid Mass Spectrometer | Thermo Fisher Scientific |  |
| Petrisoft 35 | Matrigen | #PS35-COL |
| Odyssey Sa® Infrared Imaging System | LI-COR | N/A |
| Olympus Fluoview 1000 Confocal Microscopy | Olympus | N/A |
| Vacuum concentrator | SpeedVac, Eppendorf | N/A |
| Manual Stretch Device (4 cm<sup>2</sup> uniaxial) | STREX | #ST-0040 |
| PDMS uniaxial stretch chamber (2 cm<sup>2</sup> wells) | STREX | #SC-0044 |

## MATERIALS AND EQUIPMENT

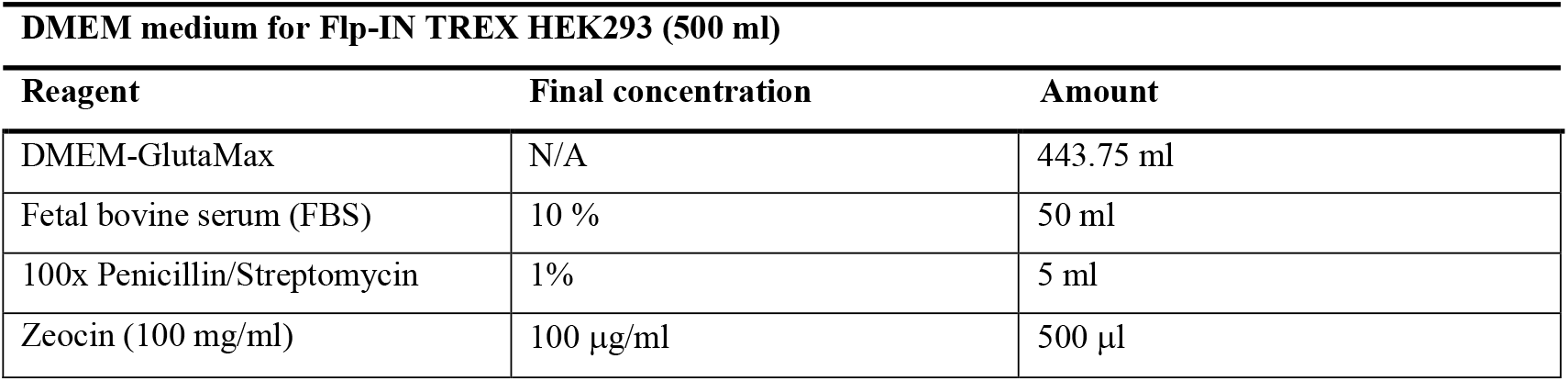
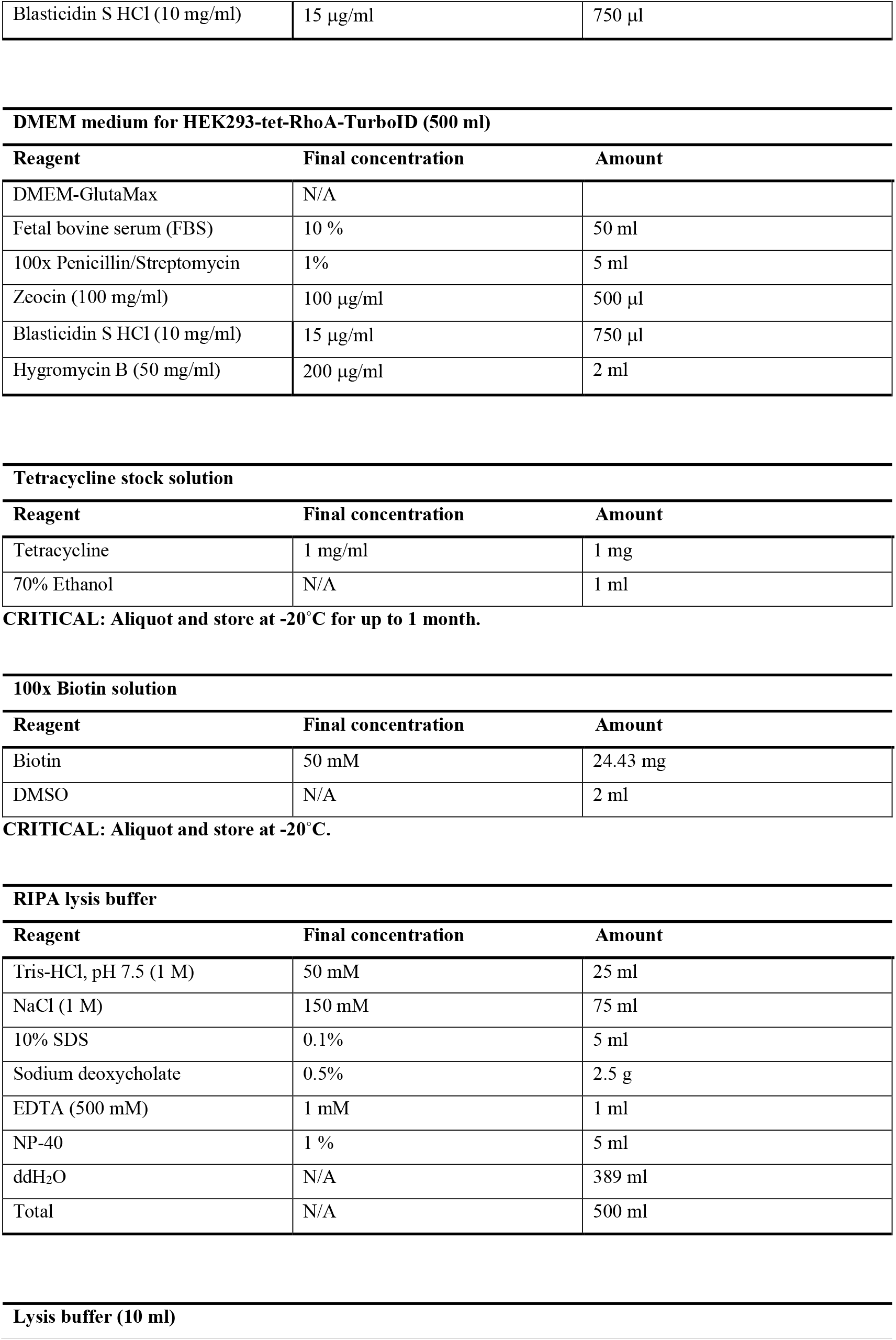
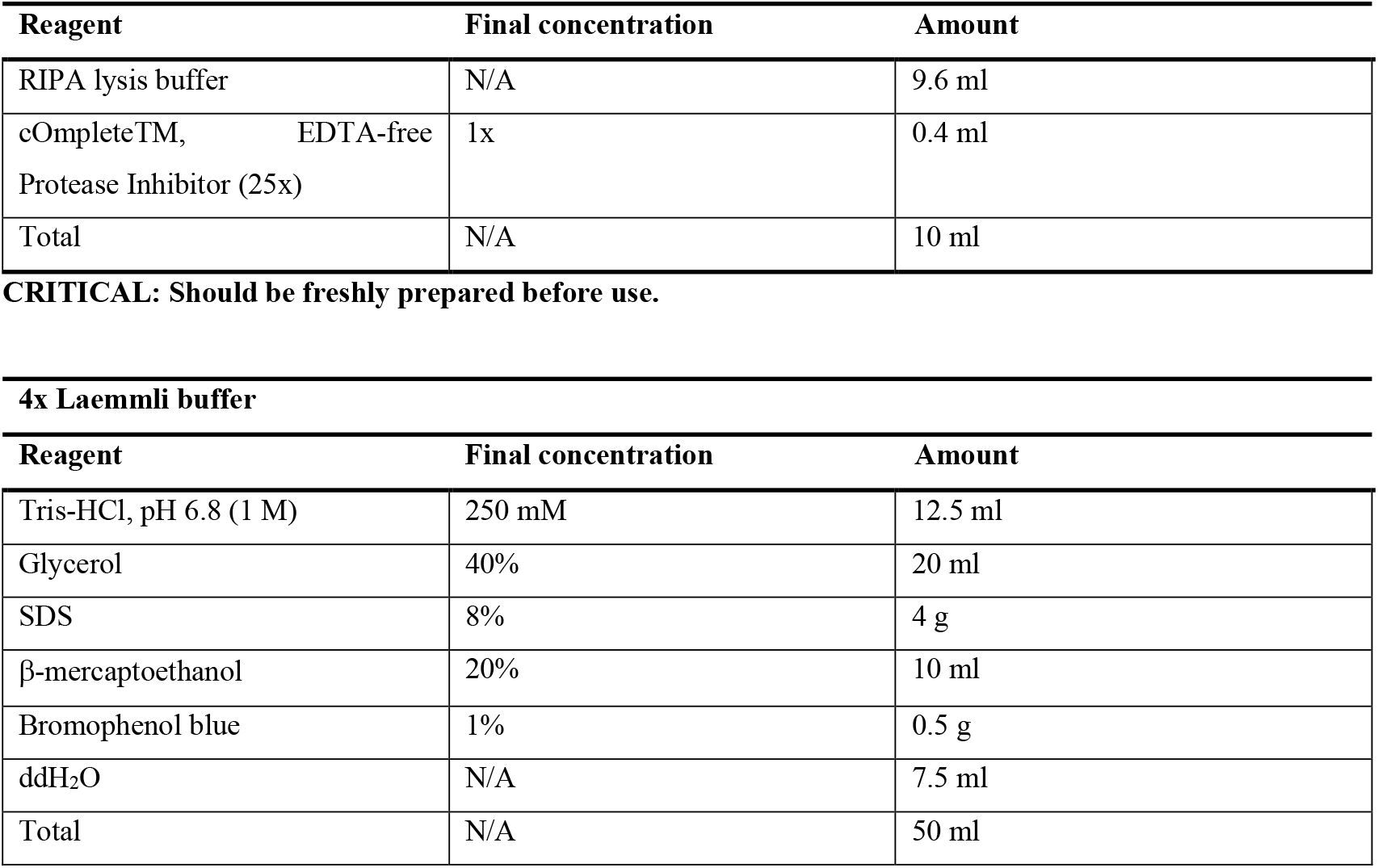

## STEP-BY-STEP METHOD DETAILS

### Generation of a cell model with tuneable actomyosin contractility

Timing: 6 weeks

This section describes details of generating a stable HEK293-tet-RhoA-TurboID cell line by sequentially introducing two plasmids described above in the before you begin section, pcDNA5/FRT/TO-RhoA-Q63L and pcDNA3.1-Puro-3xHA-TurboID-NLS, into a host cell line Flp-IN TREX HEK293. The first plasmid can integrate into the host cell genome at a FRT integration site and express constitutively active RhoA variant (RhoA-Q63L) under the control of a tetracycline inducible promoter, while the second plasmid expresses TurboID biotin ligase for proximity labelling of the nuclear proteome.

***Note*:** Before starting to generate the above cell line, a kill curve experiment to determine the optimal concentration of the selecting drug for the establishment of stable clones should be performed. In this protocol, we use 200 μg/ml hygromycin and 1.5 μg/ml puromycin to select HEK293-tet-RhoA and HEK293-tet-RhoA-TurboID stable clones respectively.

1. Generation of HEK293-tet-RhoA cell line.
  a. Seed 4×10^5^ Flp-IN TREX HEK293 cells in a 35mm culture dish, when cells have reached a confluency of about 80% (after ca. 20 h) cells are ready for transfection.
  b. Prepare a mixture of DNA containing 9:1 ratio (w/w) of pOG44: pcDNA5/FRT/TO-RhoA-Q63L.
  c. Transfect DNA mixture into Flp-IN TREX HEK293 by Lipofectamine™ 2000 according to manufacturer’s instruction.
  d. After 24 h of transfection, replace the medium with fresh complete medium containing 15 μg/ml blasticidin. Continue to culture the cells for another 24 h.

### *CRITICAL

Since the *lacZeo* gene in Flp-IN TREX HEK293 was interrupted by the integration of pcDNA5/FRT/TO-RhoA-Q63L into the FRT site, zeocin must be removed from the culture medium after transfection because the transfectants will become sensitive to zeocin.

e. Split cells (1:10) in complete medium containing 15 μg/ml blasticidin into a new 6-well culture plate. Incubate the cells for 24 h to allow cells to attach.
f. Start the selection of successful transfectants by replacing the medium with selection medium supplemented with 15 μg/ml blasticidin and 200 μg/ml hygromycin.
g. Refresh the selection medium every 2-3 days until colonies appear under the observation with an inverted microscope. Pool all colonies by detaching the cells with trypsin and replating, which is expected to result in an isogenic population given the defined integration locus for the plasmid.

### *CRITICAL

It is important to validate the generated stable HEK293-tet-RhoA cell line before introducing the second plasmid. For this, we assessed EGFP expression by live fluorescence microscopy after 24 h treatment with 0.5 μg/ml tetracycline. We expected to observe isogenic EGFP expression across cell population, confirming the stable and inducible EGFP-RhoA-Q63L expression.

2. Generation of stable HEK293-tet-RhoA-TurboID cell line

This part describes the further modification of the above cell line by introducing a nuclear localised Turbo protein biotin ligase allowing selective biotinylation of nuclear proteins.

a. Seed 3×10^5^ HEK293-tet-RhoA in a 35 mm petri dish for overnight. The cells are ready for transfection when reaching about 80% of confluency.
b. Perform stable transfection of 1 μg of linearised pcDNA3.1-Puro-3xHA-TurboID-NLS plasmid to cells using lipofectamine 2000 according to manufacturer’s instruction.
c. 24 h after transfection, replate and culture cells in a new 10 cm petri dish with complete medium for 24 h.
d. By the time cells should reach around 30∼50% confluency. Start selecting puromycin-resistant transfectants with selection medium containing 1.5 μg/ml puromycin. Change selection medium every 2∼3 days until single colonies appear.

3. Isolate single cell colonies using cloning discs.

***Note*:** All the procedure must be done under sterile conditions. Select colonies that are of average size. The Scienceware^®^ cloning disc (Z374431, Sigma-Aldrich) used in this protocol is now discontinued. However, alternative methods like using cloning cylinders can be considered to harvest the cells.

a. Examine the cells with an inverted microscope to locate single colonies that are ready for isolation. Draw a circle around the selected colonies on the bottom of the dish with a marker pen.
b. Aspirate culture medium and rinse the cells twice with prewarmed 1x PBS to remove floating cells. Leave a thin layer of PBS in the culture dish to avoid cells from drying.
c. Use sterile forceps to place cloning disc over the selected colony, press gently on the cloning disc and carefully scrape the cell colony off with the disc.
d. Transfer the disc to 24-well plate, with the disc side containing cell colony facing down.
e. Continue to culture the cells in selection medium for two days before removing the disc. It is time to remove the cloning disc when the cells have proliferated so they expanded out of the disc area.
f. When cells reach around 70% confluency, replate the cells to a suitable culture vessel for further expansion.

### Characterization of stable HEK293-tet-RhoA-TurboID cell line

Timing: 3 days

To validate the suitability of the selected clones for the intended screen, tetracycline dependent EGFP-CA-RhoA expression and protein target biotinylation should be confirmed in each stable HEK293-tet-RhoA-TurboID cell line generated (Figure 2).

**Figure 2.**
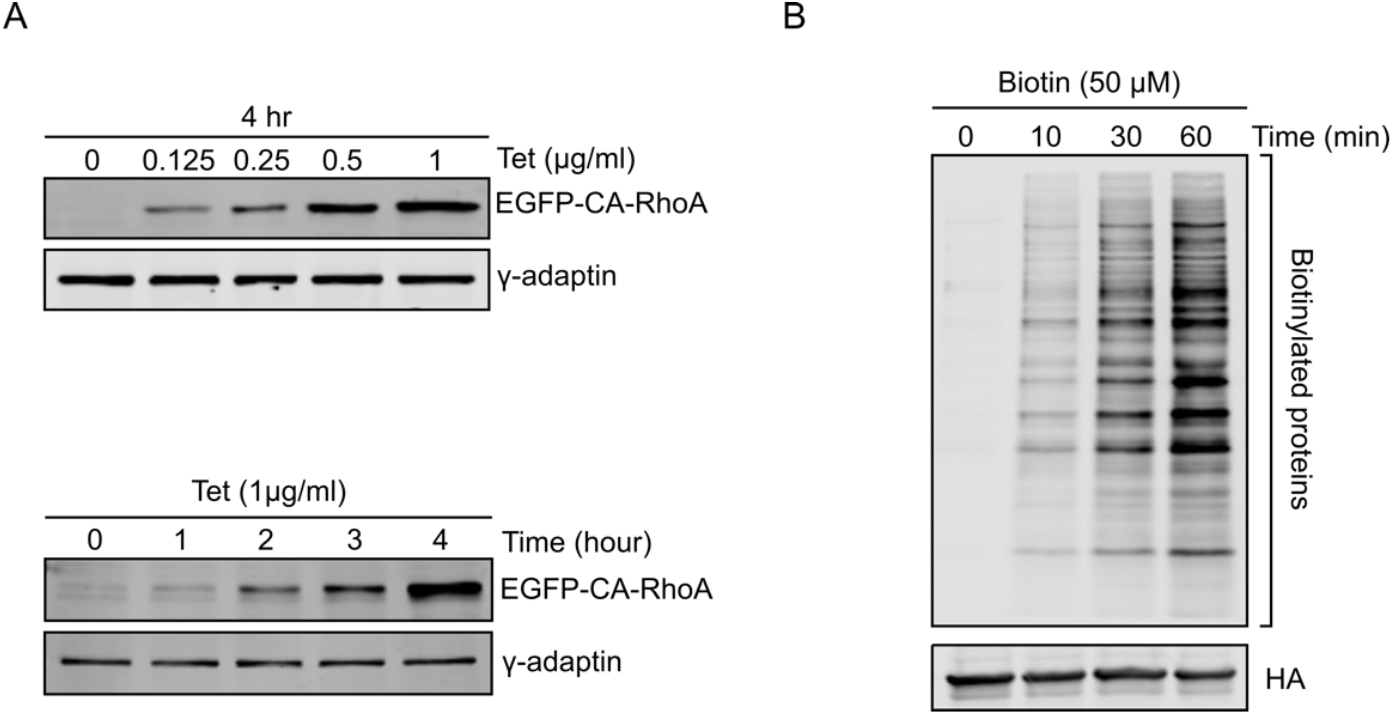
Inducible EGF-CA-RhoA and TurboID mediated biotinylation in HEK293-tet-RhoA-TurboID. **A**: EGFP-CA-RhoA expression in HEK293-tet-RhoA-TurboID can be controlled by tetracycline in a dose dependent (upper panel) and time dependent (lower panel) manner. γ-adaptin is used as loading control. **B**: TurboID-catalysed protein biotinylation in the presence of 50 μM Biotin for indicated time points. The biotinylated proteins were detected by streptavidin blotting. HA is used as loading control.

4. EGFP-CA-RhoA expression analysis.
  a. Culture 5.5×10^5^ HEK293-tet-RhoA-TurboID at high density in 35 mm culture dish for 24 h.
  b. Incubate cells with varying doses of tetracycline and for different durations at 37°C.
  c. Wash cells twice with ice-cold PBS. Prepare cell lysate using ice-cold RIPA lysis buffer following standard cell lysis procedure.
  d. Load 20 μg protein onto a 10% polyacrylamide gel and detect EGFP-CA-RhoA expression by western-blotting using EGFP antibody.
5. Protein biotinylation analysis.
  a. Culture HEK293-tet-RhoA-TurboID as Step 4a.
  b. Incubate cells with 50 μM biotin for varying time periods from 10 ∼ 60 min at 37°C.
  c. Aspirate culture medium and wash the cells five times with ice-cold PBS to remove residual biotin.
  d. Prepare cell lysate as described in Step 4c.
  e. Load 20 μg protein onto a 10% polyacrylamide gel and detect biotinylated proteins by western blotting using fluorophore-conjugated streptavidin.

### Functional validation of stable HEK293-tet-RhoA-TurboID cell line

Timing: 2 days

The stable HEK293-tet-RhoA-TurboID cell line enables tetracycline-inducible expression of CA-RhoA, which is expected to enhance actomyosin contractility and promote nuclear accumulation of YAP. To confirm this is the established HEK293-tet-RhoA-TurboID cell lines, we measure changes in cellular contractile forces and subcellular localization of YAP upon tetracycline induction (Figure 3).

**Figure 3.**
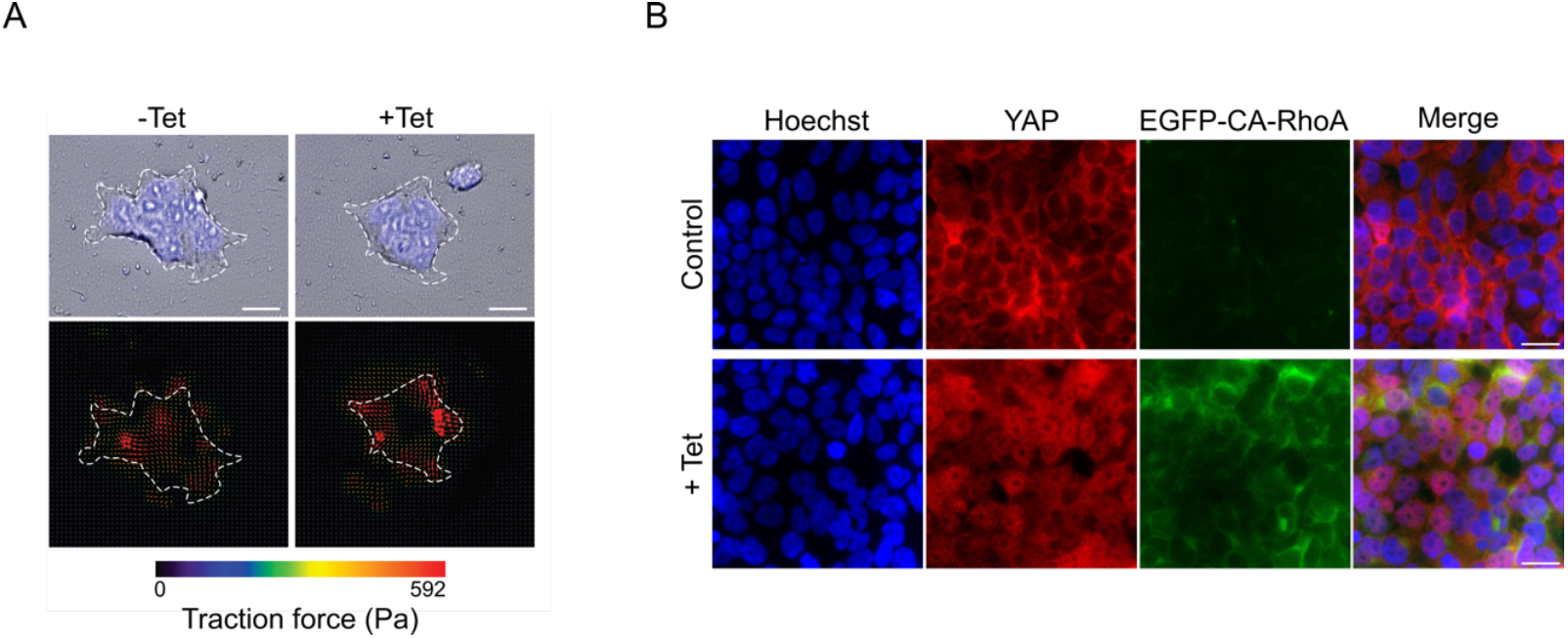
Tetracycline induced EGFP-CA-RhoA expression enhances actomyosin contractility and YAP nuclear accumulation. **A**: Detection of traction force in HEK293-tet-RhoA before and after two hours of tetracycline treatment. Cells were visualized by staining of nuclei. Red arrows indicate the force direction. Bar represents 20 μm. Data reprinted with permission from Tseng et al. [1]. **B**: Immunofluorescence imaging shows the tetracycline-induced EGFP-CA-RhoA expression and subsequent YAP nuclear accumulation. Bar represents 20 μm.

6. Assess tetracycline induced cellular contractility.
  a. Plate 1×10^5^ HEK293-tet-RhoA-TurboID on 35 mm collagen-coated plates (stiffness: 12 kPa) embedded with 0.2 µm red fluorescent beads.
  b. Incubate for 24 h under standard condition.
  c. Replace medium with fresh medium containing 0.05 µg/ml Hoechst for nucleus counterstaining.
  d. Transfer the plate to an incubation chamber (37°C, 5% CO_2_) on the microscope stage.
  e. Acquire baseline images of the cell clusters and beads using a Cell Discoverer 7 microscope (Zeiss, Germany).
  f. Treat cells with 1 µg/ml tetracycline and acquire follow-up images after 2 h (concentration and time will depend on the the concentration and time dependent EGFP-RhoA expression levels established above).
  g. Quantify beads displacement using particle image velocimetry. Calculate traction force via Fourier transform traction cytometry [17, 18].
7. Analyse subcellular localization of YAP by immunofluorescence imaging.
  a. Seed 5.5×10^5^ cells on Poly-L-Lysin coated coverslips in 12-well plate for 24 h. The cells should reach high cell density before tetracycline treatment.
  b. Treat cells with 1 μg/ml tetracycline or 0.1 μl 70% Ethanol as vehicle control for 2 h at 37°C (concentration and time will depend on the the concentration and time dependent EGFP-RhoA expression levels established above).
  c. Wash the cells three times with 1x PBS, then fix the cells with 4% PFA (500 μl/well) for 10 min at RT.
  d. Wash the cells three times with 1x PBS, then permeabilise cells with permeabilization buffer (1 ml/well) for 15 min at RT.
  e. Wash the cells three times with 1x PBS. Block with blocking buffer (1 ml/well) for 1 h at RT.
  f. Incubate with YAP primary antibody for 2 h at RT.
  g. Wash the cells three times with 1x PBST.
  h. Incubate the cells with secondary antibody and Hoechst for 1 h in the dark.
  i. Wash the cells three times with 1x PBST.
  j. Mount the coverslips onto clean glass slides with Prolong gold anti-fade mounting medium (3 μl/each slide).
  k. Analyse the subcellular localisation of YAP and EGFP-CA-RhoA expression using fluorescence microscope.

### Isolation of nuclear proteins by streptavidin pulldown

Timing: 2 days

To isolate nuclear proteins for subsequent mass spectrometry analysis, HEK293-tet-RhoA-TurboID cells are cultured at high cell density in 100 mm dishes for 24h followed by treatment with 1 μg/ml tetracycline or 1 μl 70% Ethanol as vehicle control for 2 h at 37°C to induce EGFP-CA-RhoA expression. Next, cells are labelled by addition of 500 μM biotin for 20 min at 37°C before lysis in ice-cold RIPA buffer following steps 5c-d. YAP serves as positive control given its known regulation by actomyosin contractility.

8. Aliquot 100 μl streptavidin agarose beads and wash the beads with ice-cold RIPA buffer by centrifugation at 1,800 x *g* for 2 min at 4°C. Keep the beads on ice.
9. Remove the supernatant and add the cell lysate (as prepared above) containing 3000 μg protein to the beads. Incubate overnight on a constant roller at 4°C.
10. Transfer the cell lysate/streptavidin beads to a Wizard^R^Minicolumn. Wash the beads with buffer in the following order: 10 ml of 2% SDS, 10 ml of RIPA buffer, 10 ml of 0.5M NaCl, 10 ml of 2M Urea/50 mM Tris-HCl pH8, and 10 ml of 50 mM ammonium bicarbonate.
11. Resuspend the beads in 120 μl of 50 mM ammonium bicarbonate solution, transfer the beads to a fresh Eppendorf tube.
12. Protein elution for western blot analysis.
  a. Mix 20 μl of beads from Step 11 with 20 μl of 2x Laemmli buffer.
  b. Boil for 5 min at 98 °C to elute biotinylated proteins.
13. Western blot analysis.
  a. Detect biotinylated proteins by western blotting using fluorophore-conjugated streptavidin.
  b. Analyse YAP expression in nuclear fraction by standard western blotting. Also blot the samples with nucleolin (nuclear marker) and β-tubulin (cytosolic marker) to confirm the selectivity of the streptavidin pulldown (Figure 4).

**Figure 4.**
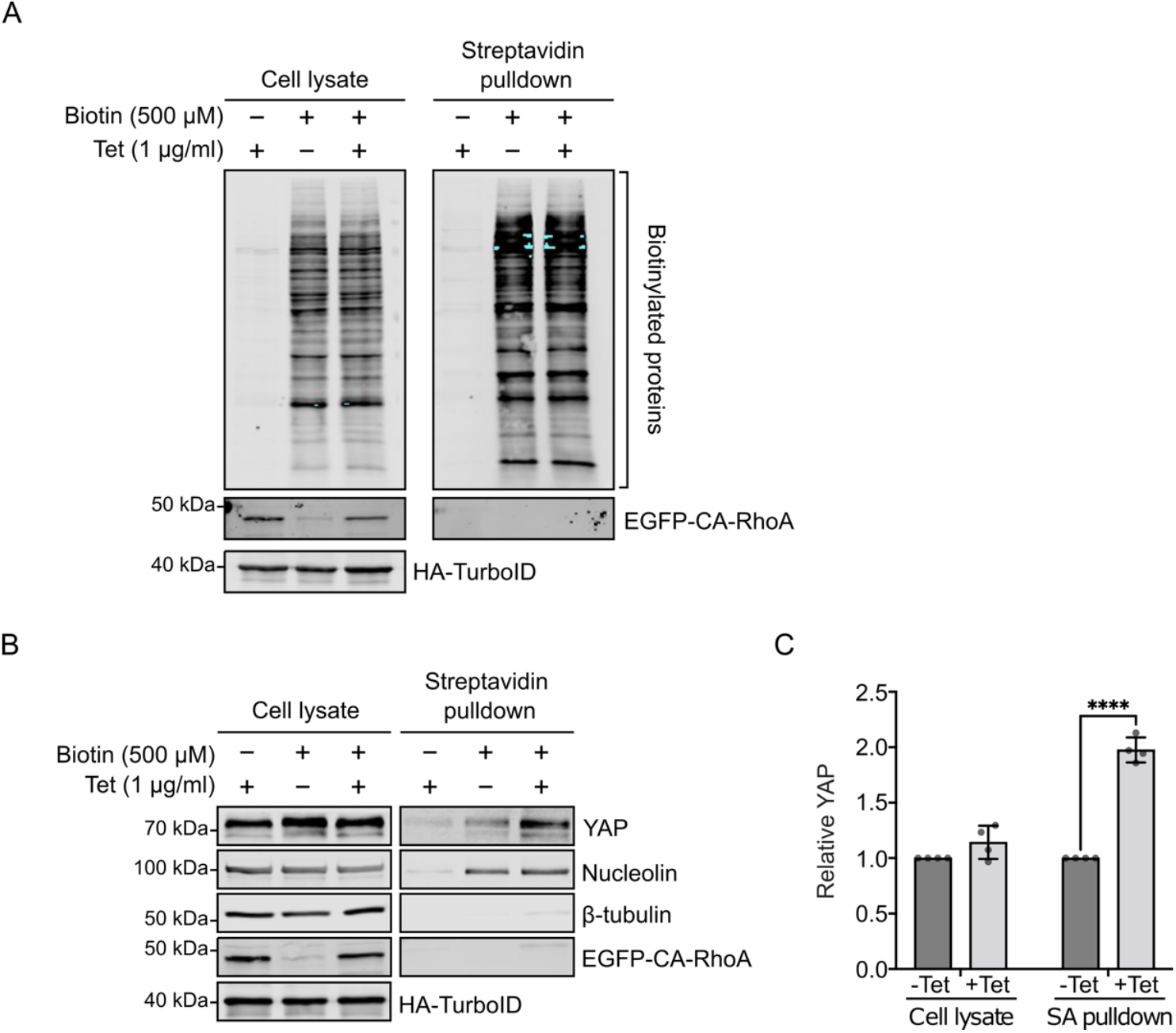
Enrichment and biotinylation of nuclear proteins in HEK293-tet-RhoA-TurboID. **A**: HEK293-tet-RhoA-TurboID were treated with tetracycline for 2 h followed by biotin for 20 min. The stably expressing TurboID was detected by western blotting with anti-HA antibody. The biotinylated proteins in samples from cell lysate and nuclear fraction (streptavidin pulldown) were analysed by western blotting with streptavidin conjugated to Alexa Fluor 680. **B**: Western blot of indicated proteins in total cell lysate and nuclear fraction (streptavidin pulldown). Nucleolin and β-tubulin were used as nuclear and cytosolic marker respectively. **C**: Quantification shows the normalised YAP expression from B. Replicates=4. Values are means ± s.d. \*\*\*\**p*<0.0001.

### Sample preparation for mass spectrometry

Timing: 2 days

This section describes the on-bead reduction, alkylation, and tryptic digestion of biotinylated proteins from Step-11 and the subsequent desalting using C18 stage tips to clean up peptide samples before mass spectrometry analysis.

14. On-bead tryptic digestion
  a. Wash the remaining cell lysate/streptavidin beads (around 100 μl) with 1 ml of 50 mM ammonium bicarbonate solution by centrifugation at 1,800 x *g* for 3 min at room temperature.
  b. Remove the supernatant. Incubate the beads in 200 μl of 50 mM ammonium bicarbonate containing 10 mM Tris(2-carboxyethyl)phosphine (TCEP) for 15 min at 37 °C with shaking.
  c. Add 4 μl of 0.5 M IAA (Iodoacetamide) to alkylate sample for 15 min in the dark at 37 °C with shaking.
  d. Add 1 μg of trypsin to digest the sample overnight at 37 °C with shaking.
  e. The following day, centrifuge sample at 1,800 x *g* for 1 min at room temperature. Transfer the supernatant (containing the peptide fragments) to a clean Eppendorf tube.
  f. Add 14 μl of 10% Trifluoroacetic acid (TFA) to acidify samples to a pH of 3. Use pH indicator strips to monitor pH changes.
15. Peptide desalting with Pierce C18 stage tip
  a. Use a clean pair of scissors to remove approximately 1/4 from the top and bottom of a 1 ml blue tip. The modified tip will connect the C18 stage tip to an uncut 1 ml tip for stable sample loading and washing.
  b. Wash the C18 stage tip with the addition of 100 μl of 0.1% TFA/50% acetonitrile (ACN). Ensure full wetting of the C18 resin for optimal binding.
  c. Transfer the acidified peptides from Step 14f into the prepared C18 stage tip assembly.
  d. Flick gently to remove air bubbles and ensure even resin contact.
  e. Slowly pipette the sample through C18 stage tip.
  f. Collect the flow-through in the original same tube. Repeat above steps for improved binding.
  g. Wash the C18 stage tip three times with 100 μl of 0.1% TFA to remove salts and impurities.
  h. Slowly elute peptides with 100 μl of 0.1% TFA/50% ACN, collecting the flow-through in a fresh 2 ml Eppendorf tube.
  i. Dry the purified peptides in a vacuum concentrator for 90 min at 45 °C.
  j. Add 10-40 μl of 0.5% formic acid and incubate at 21°C with shaking for 10 minutes. The volume used to resuspend the peptides will depend on the injection volume specified in the LC-MS/MS analysis below.
  k. Transfer the peptide solution to an autosampler vial compatible with the liquid chromatography system to be used for LC-MS/MS analysis

**Note:** C18 spin columns can be used instead of C18 Stage tips.

### Mass spectrometry data acquisition

Timing: 0.5-2 hours per sample, depending on the scan rate of the instrument and the coverage required.

This section describes the proteomic analysis using liquid chromatography tandem mass spectrometry (LC-MS/MS), peptide identification and quantification using MaxQuant and downstream data analysis using Perseus software.

16. Inject 5-10 μl of sample from Step 15j into a nanoflow liquid chromatography tandem mass spectrometry (LC-MS/MS) for peptide analysis. We performed the analysis using an Orbitrap Elite Hybrid Mass Spectrometer (Thermo Fisher Scientific) coupled online to an UltiMate RSLCnano LC System (Dionex™). The system was controlled by Xcalibur 3.0.63 (Thermo Fisher) and DCMSLink (Dionex).
  a. Run QC samples to benchmark the system performance e.g. 100 fmol of a BSA digest to check chromatography and MS sensitivity and/or 100 ng of a HeLa digest to assess proteome coverage.
  b. Generate an LC-MS/MS method that includes either a direct injection single column configuration or a two-column configuration which allows for online desalting and separation.
  c. Desalt peptides on-line using an Acclaim PepMap 100 C18 nano/capillary BioLC, 100A nanoViper 20 mm x 75 µm I.D. particle size 3 µm (Fisher Scientific) at a flow rate of 5 μl/min.
  d. Separate peptides using a 125-min gradient from 5 to 35% buffer B (0.5% formic acid in 80% acetonitrile) using an EASY-Spray column, 50 cm × 50 μm ID, PepMap C18, 2 μm particles, 100 Å pore size (Fisher Scientific) at a flow rate of 0.25 μl/min.
  e. Perform DDA LC-MS/MS analysis using an Orbitrap Elite with a cycle of one MS (in the Orbitrap) acquired at a resolution of 60,000 at m/z 400, with the top 20 most abundant multiply charged (2+ and higher) ions in a given chromatographic window subjected to MS/MS fragmentation (CID) in the linear ion trap.
  f. Set an FTMS target value of 1e6 and an ion trap MSn target value of 1e4 with the lock mass (445.120025) enabled to maintain high mass accuracy.
  g. Set the maximum FTMS scan accumulation time to 100 ms and the maximum ion trap MSn scan accumulation time to 50 ms.
  h. Enable dynamic exclusion with a repeat duration of 45 s with an exclusion list of 500 and an exclusion duration of 30.

**Note:** Sample may be analysed using either data-dependent analysis (DDA) or data-independent analysis (DIA).

**Note:** Newer generations of Orbitrap or Q-TOF instruments will offer a more sensitive and deeper analysis.

### Mass spectrometry data analysis

17. Analyse raw MS data
  a. Analyse raw data with MaxQuant (version 1.6.2.6) using an up-to-date human UniProt database (UP000005640).
  b. Set the search parameters as follows: digestion set to Trypsin/P with maximum of 2 missed cleavages, methionine oxidation and acetylation at N-terminal peptide as variable modifications, carbamidomethylation at cysteine as fixed modification,
  c. Enable between runs with a match time window of 0.7 min and a 20-min alignment time window, label-free quantification (LFQ) enabled with a minimum ratio count of 2, minimum number of neighbors of 3 and an average number of neighbors of 6.
  d. Set a first search precursor tolerance of 20 ppm and a main search precursor tolerance of 4.5 ppm for FTMS scans and a 0.5 Da tolerance for ITMS scans.
  e. Set a protein false discovery rate (FDR) of 0.01 and a peptide FDR of 0.01 for identification level cut-offs.

**Note**: Newer versions of MaxQuant are available.

**Note**: Search tolerances should be set in line with the mass accuracy of the analysers used for MS1 and MS2 scans.

18. Export MaxQuant proteinGroups output.txt file to Perseus (version 1.5.6.0) for downstream data analysis.
  a. Load the LFQ Intensities to the main category and the protein and gene names to the Text category.
  b. Group samples by giving replicates of the same group the same name using the Categorical Annotation Rows function.
  c. Log2 transform the LFQ intensity data.
  d. Remove proteins identified by fewer than three replicates in any one group.
  e. Generate a histogram of the quantitative data to assess the distribution; a normal distribution is needed for the downstream statistical analysis to be valid.
  f. To assess the reproducibility of replicates with groups, calculate the Pearson correlations using the Column correlation function and visualise using the multi-scatter plot function.
  g. Impute missing values using a downshift from the normal distribution, start with the default settings, but some adjustment may be required depending on the dataset.
  h. Perform a PCA analysis to visualise the differences between the groups and their replicates.
  i. To assess the significance of the quantitative differences between groups (e.g. +/-Tet addition to induce Rho), perform a two-sample t-test with a permutation-based FDR of 0.05. The size effect can be given weight by adjusting the S0 parameter, which is typically between 0.1 and 2.
  j. The results of the statistical analysis can be visualised by plotting a volcano plot with the same parameters.

**Note:** The adjustment of the S0 value will produce sets of significant proteins with varying fold changes, but the overall FDR will be maintained at 0.05. An additional fold-change cutoff may be applied to the data.

### Validation of mechanosensitive nuclear localization of candidate proteins

Timing: 1 week

Our screening approach for the identification of proteins with a mechanosensitve nuclear localisation is a multi-layered approach. The first layer is the unbiased identification of proteins that change nuclear levels after controlled expression of constitutive active RhoA. The second layer of our screen is to test the effect of defined mechanical cues more directly on the subcellular localisation of candidate proteins identified. The mechanical cues we have tested comprise cell density, extracellular matrix stiffness, and cell stretching. Finally, to demonstrate the direct regulation by actomyosin contractility, inhibitor experiments can be performed. We illustrate our approach using the identified splicing factor PTBP1 as an example (Figure 5).

**Figure 5.**
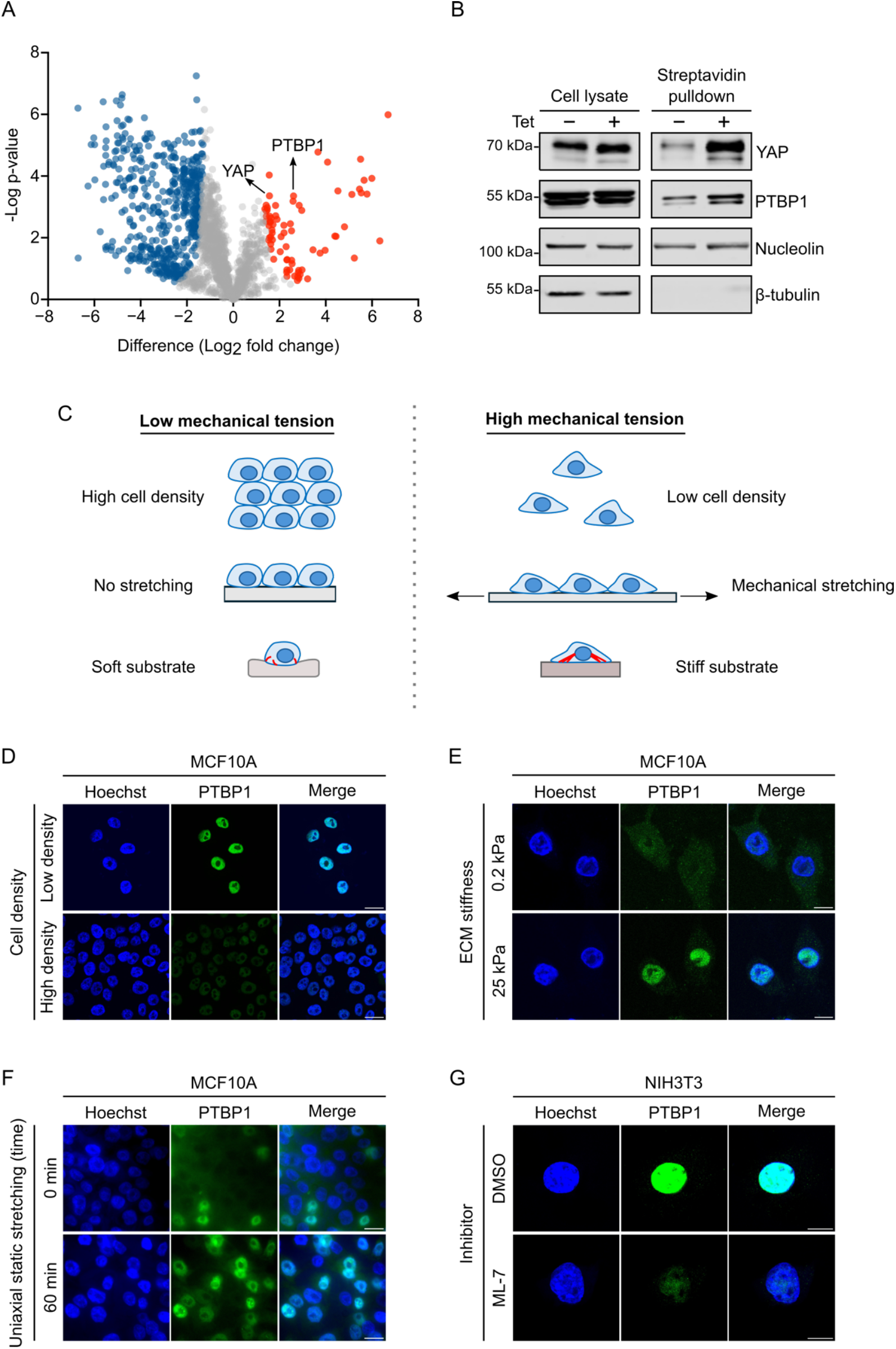
Mechanosensitive responses of PTBP1 under various mechanical conditions. **A**: Volcano plot demonstrating significant nuclear enrichment of PTBP1 following tetracycline treatment (FDR< 0.01). Reprinted data with permission from Tseng et al. [1]. **B**: Western blot analysis of PTBP1 nuclear enrichment, with YAP as a positive control. **C**: Diagram shows the mechanical conditions used in this protocol to verify PTBP1 expression. Immunofluorescence microscopy confirming the nuclear accumulation of PTBP1 induced by high mechanical tension including low cell density (**D**, scale bar: 20 μm), stiff substrate (**E**, scale bar: 10 μm), and mechanical stretching (**F**, scale bar: 50 μm). **G**: Immunofluorescence images showing PTBP1 nuclear reduction in ML-7 treated NIH3T3. Bar represents 10 μm.

19. Western blot analysis of nuclear PTBP1 levels in HEK293-tet-RhoA-TurboID.
  a. Plate 5.5×10^5^ HEK293-tet-RhoA-TurboID on 35 mm culture dish.
  b. Culture cell in puromycin-free medium for 24 h, allowing cells to high cell density.
  c. Treat the cells with 1 μg/ml tetracycline for 2 h at 37°C followed by labelling of nuclear proteins with 500 μM biotin for 20 min at 37°C.
  d. Aspirate culture medium and wash the cells five times with ice-cold PBS to remove excess biotin.
  e. Prepare cell lysate as Step 4c.
  f. Isolate and collect nuclear fraction using streptavidin pulldown as described in Step 8∼12.
20. Immunofluorescence imaging of PTBP1 in MCF10A cultured at varying cell densities.
  a. Plate MCF10A at low (0.8×10^5^) or high (5×10^5^) cell density on Poly-L-Lysine coated glass coverslips in 12-well plate.
  b. Culture the cells for 24 h under standard condition.
  c. Perform standard immunofluorescence staining as described in Step 7.
21. Immunofluorescence imaging of PTBP1 in MCF10A cultured on ECM stiffness substrate.
  a. Seed 1×10^5^ MCF10A on soft (0.2 kPa) or stiff (25 kPa) Matrigen 35 mm collagen type I coated polyacrylamide plates.
  b. Maintain culture for 3 days under standard condition, ensuring cells remain low cell density at the time of fixation.
  c. Perform standard immunofluorescence staining as described in Step 7.
  d. Analyse images using Olympus Fluoview 1000 Confocal microscopy with 60x water immersion objective.
22. Immunofluorescence staining of PTBP1 under static cell stretching. We perform uniaxial cell stretching using a manual stretch device (STREX, ST-0040) according to manufacturer’s instruction (https://strexcell.com/cell-stretching-system/#STB-100). In this protocol, we culture the cells on the 4-well PDMS uniaxial stretch chamber (2 cm^2^ wells).
  a. Prepare a 100 μg/ml collagen type I coating solution in 1 mM sterile HCl.
  b. Place the PDMS stretch chamber in a 100 mm petri dish.
  c. Add 1 ml collagen solution per well, ensure full surface coverage.
  d. Cover the petri dish with a lid and incubate the chamber at 37 °C for 4 h.
  e. Aspirate the collagen coating solution, rinse the chamber twice with serum-free culture medium.
  f. Seed 5×10^5^ MCF10A cells/well on the coated chamber and culture for 24 h under standard condition. The cells should reach high cell density before stretching.
  g. Mount the chamber on the STREX stretch device. Apply 10 % static stretch and maintain at 37 °C for 1 h.
  h. Fix the cells with 4% PFA as described in Step 7c.
  i. Aspirate PFA solution. Carefully cut out the PDMS membrane with a clean scalpel and transfer the membrane with cells up to a 35 mm dish.
  j. Perform standard immunofluorescence staining with PTBP1 as described in Step 7d∼k.
23. Immunofluorescence analysis of PTBP1 in myosin light chain kinase inhibitor ML-7 experiments.
  a. Seed 1×10^5^ NIH3T3 cells on Poly-L-Lysine coated glass coverslips in 12-well dish.
  b. Maintain the cells for 24 h under standard condition.
  c. Treat cells with 80 μM ML-7 or 0.1% DMSO (control) at 37 °C for 30 min. Effective ML-7 concentration might vary between cell types. YAP can be used as positive control to optimise concentration and incubation time.
  d. Perform standard immunofluorescence staining with PTBP1 as described in Step 7.

## EXPECTED OUTCOMES

The stable HEK293-tet-RhoA-TurboID cell line was established to modulate actomyosin contractility via controlled expression of constitutive active RhoA. This approach can be adopted also for cell lines of a different origin. If stable cell line are successfully established, cells are expected to express constitutively active RhoA variant (CA-RhoA) under the control of a tetracycline inducible promoter allowing control over time and expression levels of the EGFP tagged CA-RhoA, which can be easily monitored and quantified by western blotting (Figure 2A). The tetracycline induced CA-RhoA signalling is expected to trigger actomyosin contractility, which can be validated using traction force microscopy (Figure 3A). To further confirm mechanosensitive nuclear localisation of candidate proteins, the tetracycline induced CA-RhoA signalling should override inhibitory effect of high cell density on YAP nuclear localization with the consequence of an increase in nuclear YAP after tetracycline treatment (Figure 3B). The NLS-tagged TurboID is expected to be targeted to the nucleus to promote exclusive biotinylation of nuclear proteins in the presence of exogenous biotin, biotinylation can be monitored via western-blotting using conjugated streptavidin (Figure 4A). To confirm the specificity of the labelling for nuclear proteins the streptavidin pulldown fraction should contain the nuclear marker protein nucleolin but should be devoid of cytosolic marker proteins such as β-tubulin. YAP should be included as a positive control and should be enriched in the pulldown samples treated with tetracycline compared to mock treated samples (Figure 4B, C).

For the downstream proteomics analysis, we applied a threshold of a 2-fold change. This analysis identified 74 proteins enriched in the nucleus after tetracycline treatment. YAP, served as positive control, and should be found among the enriched protein hits to validate the screen approach (Figure 5A). The results from the proteomic approach should be further validated via western blotting (Figure 5B). In order to confirm that the nuclear accumulation of candidate proteins is due to changes in actomyosin contractility, further experiments are needed. Proteins are expected to demonstrate changes in nuclear accumulation in response to ML-7 inhibitor treatment affecting actomyosin contractility or in response to mechanical cues known to affect actomyosin contractility such as cell density, cell size, ECM stiffness or cell stretching [19-22] (Figure 5C∼G).

## LIMITATIONS

Our screening approach is based on the assumption that mechanical forces applied to the nucleus alter nucleocytoplasmic shuttling of proteins by affecting nuclear pore permeability and transport dynamics. However, given the two-hour timescale of tetracycline treatment, we cannot entirely exclude the possibility that some observed changes in nuclear protein levels result from rapid transcription and translation, reflecting mechanosensitive gene expression. Importantly, this limitation does not undermine the potential involvement of these candidate proteins in a mechanotransduction pathway.

Furthermore, RhoA/ROCK signalling can also be activated by mechanical cue-independent pathways, primarily through receptor signalling, and as part of cellular stress responses [23-25]. Thus, several of the nuclear enriched proteins found in the mass spectrometry screen might reflect mechanical cue independent RhoA/ROCK signalling and thus follow up validation using more direct mechanical cues as described in our protocol above is essential [24, 26].

## TROUBLESHOOTING

### Problem 1

Unhealthy cells caused by excessive CA-RhoA mediated actomyosin contractility. Potential solution:

Optimize both tetracycline dose and induction time. Monitor CA-RhoA expression by western blot. Evaluate cell morphology of each tetracycline treatment condition using microscope. Select the optimal concentration and duration that induce YAP nuclear enrichment while remaining normal cell viability and morphology.

### Problem 2

Unexpected nuclear YAP caused by the selection antibiotic Puromycin (Figure 6A).

**Figure 6.**
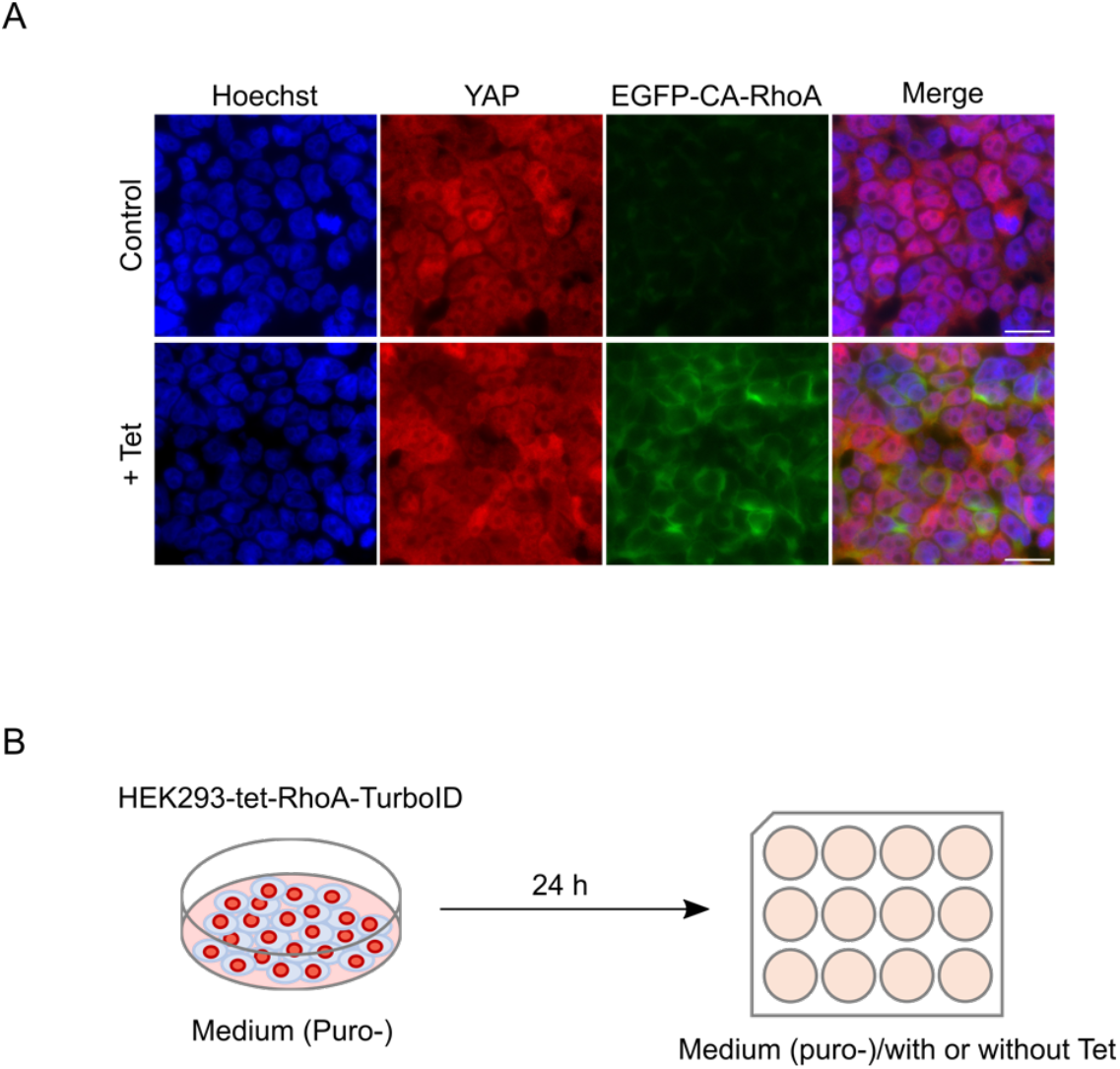
Persistent nuclear localization of YAP in HEK293-tet-RhoA-TurboID. **A**: Immunofluorescence images showing nuclear accumulation of YAP in HEK293-tet-RhoA-TurboID, independent of tetracycline treatment. Scale bar: 20 μm. **B**: Schematic diagram illustrating the preculture of cells in puromycin-free medium for 24 h prior to induction of EGFP-CA-RhoA expression by tetracycline.

### Potential solution

Only use medium supplemented with puromycin during standard cell culture to maintain genetic integrity of the stable HEK293-tet-RhoA-TurboID line. Replenish culture medium without puromycin at least 24 hr prior to tetracycline treatment and biotin labelling (Figure 6B).

### Problem 3

Heterogenous biotinylation during proximity labelling in high density culture, may reduce nuclear proteome coverage and induce background noise.

### Potential solution

Optimize both biotin concentration and treatment duration to balance labelling efficiency while minimizing the background. After labelling, thoroughly wash cells five times with ice-cold PBS to remove unincorporated biotin. Use proper amounts of streptavidin agarose beads to recover biotinylated proteins while minimize non-specific binding. For sample preparation for mass spectrometry analysis, include YAP as positive control for all streptavidin pulldown experiments to minimize batch-to-batch variation.

## LEAD CONTACT

Further information and requests of resources and reagents should be directed to by the lead contact, Kai Erdmann

## MATERIALS AVAILABILITY

This study did not genereate new unique reagents.

## DATA AND CODE AVAILABILITY

Proteomic data have been published and deposited (Tseng et al. 2025)

## SUPPLEMENTAL INFORMATION

None

## ACKNOWLEDGEMENTS

The plasmid pcDNA3-EGFP-RhoA-Q63L was gift from Gary Bokoch (Addgene plasmid # 12968), and 3xHA-TurboID-NLS_pCDNA3 was a gift from Alice Ting (Addgene plasmid # 107171). We thank Annica K. B. Gad, Ahmed Salem, and <u>Sarah C Macfarlane</u> for traction force microscopy measurements. We also like to thank the Wolfson Light Microscopy Facility, University of Sheffield, for their expert assistance with Olympus FV1000 confocal imaging. This work was supported by a fee scholarship and a publication scholarship from the University of Sheffield awarded to Weiwei Sun, and by a self-funded PhD undertaken by Pei-Li Tseng.

## AUTHOR CONTRIBUTIONS

All authors contributed to the writing and editing of the manuscript. P.-T. and W.S. performed the unpublished experiments. M.O.C. and P.-L.T. performed the mass spectrometry analysis. K.E., M.O.C, and P.-L.T. conceptualised the screening approach.

## REFERENCE

1. Tseng, P.-L., et al., Mechanical control of the alternative splicing factor PTBP1 regulates extracellular matrix stiffness induced proliferation and cell spreading. iScience, 2025. 28(4): p. 112273.

2. Jaalouk, D.E. and J. Lammerding, Mechanotransduction gone awry. Nat Rev Mol Cell Biol, 2009. 10(1): p. 63–73.

3. Kuehlmann, B., et al., Mechanotransduction in Wound Healing and Fibrosis. Journal of Clinical Medicine, 2020. 9(5): p. 1423.

4. Martino, F., et al., Cellular Mechanotransduction: From Tension to Function. Frontiers in Physiology, 2018. Volume 9 - 2018.

5. Ohashi, K., S. Fujiwara, and K. Mizuno, Roles of the cytoskeleton, cell adhesion and rho signalling in mechanosensing and mechanotransduction. The Journal of Biochemistry, 2017. 161(3): p. 245–254.

6. Burridge, K., E. Monaghan-Benson, and D.M. Graham, Mechanotransduction: from the cell surface to the nucleus via RhoA. Philos Trans R Soc Lond B Biol Sci, 2019. 374(1779): p. 20180229.

7. O’Connor, K. and M. Chen, Dynamic functions of RhoA in tumor cell migration and invasion. Small GTPases, 2013. 4(3): p. 141–7.

8. Lombardi, M.L., et al., The Interaction between Nesprins and Sun Proteins at the Nuclear Envelope Is Critical for Force Transmission between the Nucleus and Cytoskeleton*. Journal of Biological Chemistry, 2011. 286(30): p. 26743–26753.

9. Miroshnikova, Y.A. and S.A. Wickström, Mechanical Forces in Nuclear Organization. Cold Spring Harb Perspect Biol, 2022. 14(1).

10. Elosegui-Artola, A., et al., Force Triggers YAP Nuclear Entry by Regulating Transport across Nuclear Pores. Cell, 2017. 171(6): p. 1397-1410.e14.

11. Maurer, M. and J. Lammerding, The Driving Force: Nuclear Mechanotransduction in Cellular Function, Fate, and Disease. Annu Rev Biomed Eng, 2019. 21: p. 443–468.

12. Thomas, C.H., et al., Engineering gene expression and protein synthesis by modulation of nuclear shape. Proceedings of the National Academy of Sciences, 2002. 99(4): p. 1972–1977.

13. Dupont, S., et al., Role of YAP/TAZ in mechanotransduction. Nature, 2011. 474(7350): p. 179–83.

14. Das, A., et al., YAP Nuclear Localization in the Absence of Cell-Cell Contact Is Mediated by a Filamentous Actin-dependent, Myosin II- and Phospho-YAP-independent Pathway during Extracellular Matrix Mechanosensing *. Journal of Biological Chemistry, 2016. 291(12): p. 6096–6110.

15. Branon, T.C., et al., Efficient proximity labeling in living cells and organisms with TurboID. Nat Biotechnol, 2018. 36(9): p. 880–887.

16. May, D.G., et al., Comparative Application of BioID and TurboID for Protein-Proximity Biotinylation. Cells, 2020. 9(5): p. 1070.

17. Martiel, J.L., et al., Measurement of cell traction forces with ImageJ. Methods Cell Biol, 2015. 125: p. 269–87.

18. Lee, S. and S. Kumar, Cofilin is required for polarization of tension in stress fiber networks during migration. J Cell Sci, 2020. 133(13).

19. Hale, C.M., S.X. Sun, and D. Wirtz, Resolving the Role of Actoymyosin Contractility in Cell Microrheology. PLOS ONE, 2009. 4(9): p. e7054.

20. Dasgupta, I. and D. McCollum, Control of cellular responses to mechanical cues through YAP/TAZ regulation. Journal of Biological Chemistry, 2019. 294(46): p. 17693–17706.

21. Humphrey, J.D., E.R. Dufresne, and M.A. Schwartz, Mechanotransduction and extracellular matrix homeostasis. Nat Rev Mol Cell Biol, 2014. 15(12): p. 802–12.

22. Sethi, K., E.J. Cram, and R. Zaidel-Bar, Stretch-induced actomyosin contraction in epithelial tubes: Mechanotransduction pathways for tubular homeostasis. Semin Cell Dev Biol, 2017. 71: p. 146–152.

23. Mong, P.Y., et al., Activation of Rho kinase by TNF-alpha is required for JNK activation in human pulmonary microvascular endothelial cells. J Immunol, 2008. 180(1): p. 550–8.

24. Yu, O.M. and J.H. Brown, G Protein-Coupled Receptor and RhoA-Stimulated Transcriptional Responses: Links to Inflammation, Differentiation, and Cell Proliferation. Mol Pharmacol, 2015. 88(1): p. 171–80.

25. Aghajanian, A., et al., Direct Activation of RhoA by Reactive Oxygen Species Requires a Redox-Sensitive Motif. PLOS ONE, 2009. 4(11): p. e8045.

26. Tabuchi, A., et al., Nuclear translocation of the SRF co-activator MAL in cortical neurons: role of RhoA signalling. Journal of Neurochemistry, 2005. 94(1): p. 169–180.

